# Molecular signatures associated with ZIKV exposure in human cortical neural progenitors

**DOI:** 10.1101/071183

**Authors:** Feiran Zhang, Christy Hammack, Sarah C. Ogden, Yichen Cheng, Emily M. Lee, Zhexing Wen, Xuyu Qian, Ha Nam Nguyen, Yujing Li, Bing Yao, Miao Xu, Tianlei Xu, Li Chen, Zhiqin Wang, Hao Feng, Wei-Kai Huang, Ki-jun Yoon, Chao Shan, Luoxiu Huang, Zhaohui Qin, Kimberly M. Christian, Pei-Yong Shi, Mingjiang Xu, Menghang Xia, Wei Zheng, Hao Wu, Hongjun Song, Hengli Tang, Guo-Li Ming, Peng Jin

## Abstract

Zika virus (ZIKV) infection causes microcephaly and has been linked to other brain abnormalities. How ZIKV impairs brain development and function is unclear. Here we systematically profiled transcriptomes of human neural progenitor cells exposed to Asian ZIKV^C^, African ZIKV^M^, and dengue virus (DENV). In contrast to the robust global transcriptome changes induced by DENV, ZIKV has a more selective and larger impact on expression of genes involved in DNA replication and repair. While overall expression profiles are similar, ZIKV^C^, but not ZIKV^M^, induces upregulation of viral response genes and TP53. P53 inhibitors can block the apoptosis induced by both ZIKV^C^ and ZIKV^M^ in hNPCs, with higher potency against ZIKV^C^-induced apoptosis. Our analyses reveal virus- and strain-specific molecular signatures associated with ZIKV infection. These datasets will help to investigate ZIKV-host interactions and identify neurovirulence determinants of ZIKV.

## INTRODUCTION

Zika virus (ZIKV), a mosquito-borne flavivirus discovered in 1947, was responsible for only sporadic cases of infection throughout Africa and Asia until the 2007 Micronesia and 2013 French Polynesia outbreaks (1). Currently, a large-scale ZIKV outbreak is occurring in the Americas and the virus has so far spread to over 60 different countries and territories. Increasing reports link recent ZIKV infections to various forms of neuropathology, including its causal role in disorders of fetal brain development and its association with Guillain-Barré syndrome (GBS) (2). Several types of mosquitoes are capable of transmitting ZIKV to humans including the *Aedes aegypti*, which is broadly distributed throughout the world. Instances of sexual transmission have also been reported (3–5), and vertical transmission from infected pregnant women to fetuses is highly likely (6,7), albeit via an unknown mechanism (8).

ZIKV was first isolated from a sentinel *Rhesus* monkey in the Zika forest, Uganda in 1947. Akin to its close relatives in the *Flaviviridae* family, such as dengue (DENV), yellow fever, Japanese encephalitis, and West Nile viruses, ZIKV has an icosahedral outer envelope and a dense inner core containing one single-strand positive-sense RNA genome between 10 and 11 kb in length (9–12). Of the two distinct lineages of ZIKV (African and Asian), the strains currently circulating in the Western Hemisphere are more closely related to the Asian lineage than to the African lineage (13). In previous outbreaks, an estimated 80% of ZIKV-infected individuals were asymptomatic and the rest showed only mild symptoms (data from the Centers for Disease Control and Prevention (CDC), Atlanta, GA, USA). In contrast, one striking feature of the current ZIKV epidemic is the association of viral infection with an increased risk of congenital microcephaly and serious neurologic complications, such as GBS in adults (14). This increased risk of congenital microcephaly following ZIKV infection appears to be rare among flaviviruses. For example, despite similarities in protein sequences and insect vectors, DENV has not been linked to the congenital microcephaly associated with ZIKV. Consistent with mosquitoes being the primary transmission route, dermal fibroblasts, epidermal keratinocytes, and immature dendritic cells were found to be permissive for ZIKV infection (15). Furthermore, ZIKV of Asian origin was present in the amniotic fluid of two pregnant Brazilian women diagnosed with fetal microcephaly (16), supporting the notion that ZIKV can pass the placental barrier. ZIKV RNA has also been detected in various organs of fetuses with microcephaly, with the highest viral loads found in fetal brain tissue (7,17).

To establish a direct link between ZIKV and microcephaly, we and others have shown that ZIKV efficiently infects human neural progenitor cells (hNPCs) in monolayer and three-dimensional organoids derived from induced pluripotent stem cells (18–21) and that its efficiency in infecting neurons, induced pluripotent stem cells (iPSCs) and human embryonic stem cells (hESCs) is much lower (18). Infected hNPCs further release infectious ZIKV particles. Importantly, ZIKV infection increases cell death and dysregulates cell-cycle progression, resulting in reduced proliferation of forebrain-specific hNPCs and reduced neuronal layer thickness in cerebral organoids, supporting a direct link between ZIKV infection and cortical development. In one recently reported clinical case (7), postmortem analysis revealed diffuse cerebral cortex thinning in a fetal brain infected by the ZIKV strain of the Asian genotype. This experimental evidence, along with the epidemiological correlation and clinical isolation data, supports the conclusion that ZIKV plays a causal role in microcephaly (2). How ZIKV could specifically impair brain development and functions remains to be determined.

Here we systematically profiled the transcriptomes of hNPCs derived from human induced pluripotent stem cells (hiPSCs) upon exposure to ZIKV^M^ (MR766 strain, African lineage), ZIKV^C^ (FSS13025 Cambodian strain, Asian lineage), or DENV (Thailand isolate 16681, serotype 2), and compared the gene expression changes among different strains and viruses. Our analyses reveal virus- and strain-specific molecular signatures associated with ZIKV infection. Datasets presented here could be an important resource for understanding the molecular pathogenesis of ZIKV-induced abnormalities in fetal and adult brain, and for developing effective therapeutic approaches to combat ZIKV infections and its consequences.

## MATERIALS AND METHODS

### Culture of human iPSCs and differentiation into cortical neural progenitor cells

Human iPSCs were cultured and differentiated into cortical neural progenitor cells as described previously (22–25). A full description is available in Supplementary Data.

### Preparation of viruses

ZIKV^M^ (strain MR766) was used to infect Aedes C6/36 mosquito cells at an MOI of 0.02. Supernatant was collected on day 6 post-infection, filtered, and stored at −80°C for titering and infection studies. ZIKV^C^ (strain FSS13025) was obtained from World Reference Center for Emerging Viruses and Arboviruses (WRCEVA), produced in C6/36 cells, and titered on Vero cells. The virus stock was stored at −80°C for titering and infection studies. DENV (DENV type 2, strain 16681) was obtained from BEI Resources and used to infect Aedes C6/36 mosquito cells at an MOI of 0.05. Supernatant was collected on day 6 post-infection, filtered and stored at −80°C for titering and infection studies. Virus stocks were titered on Vero cells using an immunostaining-based focus-forming-units (ffu) assay. An equal volume of supernatant from uninfected C6/36 cells was collected for mock infection.

### Cell infection

DENV and ZIKV^M^ infections on hNPCs were performed at different times. One million hNPCs were seeded into 12-well plates one day before infection. Virus was added to cells at a reduced volume for a 2-hr incubation, at an MOI of 0.08, followed by a PBS wash and the addition of fresh medium. The infection duration was 67 hr for DENV, and 56 hr for ZIKV^M^. The time points were optimized for each virus to maximize infection rate while minimizing cell death. For ZIKV^M^ and ZIKV^C^ infections on hNPCs, hNPCs were seeded into T-25 flasks. Virus was added to cells at a reduced volume for a 2-hr incubation, followed by a PBS wash and the addition of fresh medium. The MOI was 0.02 for ZIKV^M^. To achieve similar infection rate, an MOI of 0.04 was used for ZIKV^C^. The infection duration was 64 hr for both strains. Cells were harvested by gentle scraping in ice-cold PBS. Cells from three wells were combined per biological replicate (total of approximately 420,000 cells per unique sample).

### Immunocytochemistry

Cells were fixed with 4% paraformaldehyde (Sigma) for 15 min at room temperature. Samples were permeabilized and blocked with 0.25% Triton X-100 (Sigma) and 10% donkey serum in PBS for 20 min as previously described (22,24,25). A full description is available in Supplementary Data.

### RNA isolation, RNA-seq library preparation, and sequencing

Total cellular RNA was purified from cell pellets using the TRIzol Reagent (Invitrogen) according to the manufacturer’s instructions. RNA-seq libraries were generated from 1 μg of total RNA from duplicated or triplicated samples per condition using the TruSeq LT RNA Library Preparation Kit v2 (Illumina) following the manufacturer’s protocol. A full description is available in Supplementary Data.

### Bioinformatic analyses

PE RNA-seq reads were first aligned to human transcriptome annotations and genome assembly (hg19) using TopHat v2.0.13 (26,27). If there were SR reads generated for the same library, the 100-cycle SR reads were then aligned using TopHat with supplementary raw junctions obtained from the first run for PE reads. The numbers of mapped reads for each sample can be found in Table S1 and S2. FPKM (<u>f</u>ragments <u>p</u>er <u>k</u>ilobase of transcript per <u>m</u>illion mapped reads) values were calculated by Cufflinks v2.2.1 (28). All data analyses were performed using R/ Bioconductor programming language. Reads mapped to the bodies of RefSeq genes were obtained using Bioconductor (29). Numbers of reads mapped to each gene were used to represent the gene expression values. Pairwise comparisons between infected and mock conditions were performed to detect differentially expressed (DE) genes using the Bioconductor package DSS (30). DSS uses a negative binomial (NB) model, which is a Gamma-Poisson compound distribution, to capture the biological and sampling variations in the expression counts. A shrinkage estimator based on Bayesian hierarchical model is implemented to combine information from all genes to improve the estimation of the dispersion parameter in NB distribution, which represents the biological variance. Hypothesis testing for differential expression was achieved by a Wald test, and the estimation of false discovery rate (FDR) was performed through the local FDR procedure (31). We have shown that DSS performs favorably compared to other methods developed for read counts, especially when sample size is small or biological variation is large (30). DE genes are defined as the ones with FDR less than 0.05. Gene ontology (GO) analyses on biological process, cellular compartment, and molecular function were performed by the Database for Annotation, Visualization and Integrated Discovery (DAVID) v6.7 (32). To identify significantly enriched GO terms using DAVID, a Benjamini–Hochberg procedure was used to control FDR at 0.05. Pathway enrichment, and protein interaction network nodule analyses were performed by the WEB-based Gene SeT AnaLysis Toolkit (WebGestalt) update 2015 (33) with adjusted p-value no bigger than 0.05 and by GeneMANIA (34).

### Statistics

Student’s *t*-tests and Chi-square tests were performed by Prism 6 (GraphPad Software).

### Data access

RNA-seq data reported in this paper have been submitted to Gene Expression Omnibus (http://www.ncbi.nlm.nih.gov/geo/) with accession number GSE80434.

## RESULTS

### Human neural progenitor cells are permissive to DENV

Both DENV and ZIKV belong to the *Flavivirus* genus of the *Flaviviridae* family. These two flaviviruses are primarily transmitted by the same mosquito vectors, and the amino acid sequences of their polyproteins share a high degree of similarity (55.6% identical, 81.0% similar, DENV2 vs. ZIKV^M^). To determine whether DENV, like ZIKV, can infect hiPSC-derived hNPCs, we exposed forebrain-specific hNPCs differentiated from hiPSCs to the serotype 2 DENV (DENV2) isolate 16681, which has been widely used for DENV pathogenesis research (35-38). DENV2 efficiently infected hNPCs in vitro (Figure 1A), resulting in 84.2 ± 6.7 % (n = 6) infection of cultures at 67 hours post-inoculation with a multiplicity of infection (MOI) of 0.08, This infection efficiency was comparable to that of ZIKV^M^ in these cells under similar MOI and infection duration, as we published recently (18).

**Figure 1.**
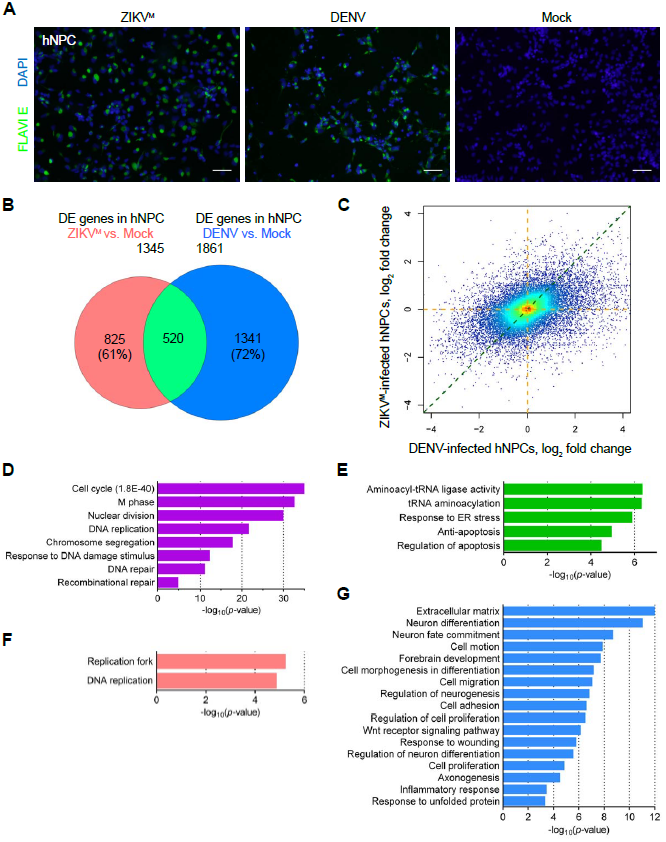
Distinct gene expression changes caused by ZIKV and DENV in hNPCs. (**A**) Sample images of immunostaining of forebrain-specific hNPCs 56 hours after infection with ZIKV^M^ and 67 hours after infection with DENV at MOIs of 0.08. Flavivirus envelop protein (FLAVIE; green)-positive cells were infected by flavivirus. DAPI (blue) was used to stain nuclei. Scale bars, 50 μm. (**B**) Weighted 2-D Venn diagram showing overlap (pale green) of differentially expressed (DE) genes identified by RNA-seq from ZIKV^M^-infected hNPCs at 56 hours post-infection (salmon) and DENV-infected hNPCs at 67 hours postinfection (royal blue) (both MOIs = 0.08). (**C**) Scatter plot of global differences in transcriptome changes between ZIKV^M^- and DENV-infected hNPCs. Log_2_ fold change of FPKM (<u>f</u>ragments <u>p</u>er <u>k</u>ilobase of transcript per <u>m</u>illion mapped reads) values from RNA-seq data are plotted. Regions with higher density of genes are shown in warmer colors. (**D**, **E**, **F**, and **G**) Gene Ontology (GO) analyses showing biological pathways and molecular functions enriched in specific sets of genes from virus-infected hNPCs. Colored magenta, genes significantly downregulated in both ZIKV^M^- and DENV-infected cells (D); lime green, genes significantly upregulated in both ZIKV^M^- and DENV-infected cells (E); salmon, genes significantly altered only in ZIKV^M^-infected hNPCs (F); and royal blue, gene significantly altered only in DENV-infected hNPCs (G). See also Figure S1 and Table S1.

### DENV and ZIKV infection causes distinct gene expression changes in hNPCs

To investigate whether ZIKV has a different functional impact on hNPCs at the molecular level, global transcriptome analyses (RNA-seq) were performed using hNPCs infected with DENV or ZIKV^M^ with similar MOIs (∼0.08) and harvested at 67 and 56 hours post-infection, respectively. Systematic analyses were performed using a DSS algorithm that we developed previously, which utilizes a negative binomial model to capture biological and sample variability in the expression counts, and a shrinkage estimator to combine information from all genes to improve the estimation of biological variance (30). We identified a large number of differentially expressed genes (DEGs) upon infection with either virus (Figure S1 and Table S1). Comparative analyses revealed distinct profiles of gene expression changes caused by DENV and ZIKV^M^ infection (Figure 1B). A total of 520 genes, representing 28% and 39% of DEGs induced by DENV and ZIKV^M^ respectively, were shared between these two sets. In addition, transcriptome-wide analysis of fold changes in gene expression indicated that, under our experimental conditions, DENV-infected hNPCs not only contain more DEGs, but also exhibit more robust transcriptome changes than ZIKV^M^-infected hNPCs (Figure 1C). For genes that were differentially expressed in at least one infection group, 74.2% of them had on average 2.91-fold larger absolute fold changes of expression in DENV-infected cells than in ZIKV^M^-infected cells, while only 25.8% of them had on average 1.66-fold larger absolute fold changes of expression in ZIKV^M^-infected cells than in DENV-infected cells (Table S1). Together these data suggest that DENV and ZIKV^M^ infections induce distinct gene expression changes in hNPCs.

### Virus-specific differentially expressed genes are enriched in DNA replication/repair, antiviral and developmental signaling pathways

To further understand the impact of DENV and ZIKV^M^ infections on the biological processes in hNPCs, we used gene ontology (GO) analyses to predict potential signaling pathways and cellular processes that were altered upon viral infection. We found a particular enrichment for 78, 32 and 26 genes related to “cell cycle”, “DNA replication” and “DNA repair”-pathways that were downregulated in both DENV- and ZIKV^M^-infected hNPCs, respectively (Figure 1D). We also observed a noticeable enrichment of genes related to protein synthesis and apoptosis-related pathways that were upregulated in both groups (Figure 1E). For virus-specific transcriptional regulation, we identified 19 and 8 genes involved in “DNA replication” and “replication fork” GO terms, respectively, and downregulated only in ZIKV^M^-infected hNPCs, but not in DENV-infected hNPCs (Figure 1F and 2A). For example, the expression of DNA2 (DNA replication helicase/nuclease 2, a protein involved in the maintenance of DNA stability) was downregulated by 64.7% upon ZIKV^M^ infection (p = 0.0057), but was decreased by only 8.6% upon DENV infection (p = 1) according to RNA-seq data. Downregulated expression of DNA2, OGG1, PRIM1, RFC4, RBM14, and RBBP7 in ZIKV^M^-infected hNPCs were further validated by quantitative RT-PCR (qRT-PCR) (Figure 2C). In contrast, among the genes specific to DENV infection that were either upregulated (665 genes) or downregulated (676 genes) in DENV-infected cells but not changed in ZIKV^M^-infected cells, none were significantly related to these three GO terms associated with DNA replication and repair. Interestingly, DENV-specific DEGs, primarily the downregulated ones, were enriched in neuron differentiation, brain development, and Wnt signaling-related pathways (Figure 1G and 2D). Furthermore, DENV-specific DEGs were also enriched in inflammatory response-related pathways, with 24 upregulated and 14 downregulated genes (Figure 1G and 2B). Therefore, DENV and ZIKV^M^ infections altered biological processes in hNPCs distinctly, with ZIKV^M^ exhibiting a broader impact on the expression of genes involved in DNA replication and DNA repair.

**Figure 2.**
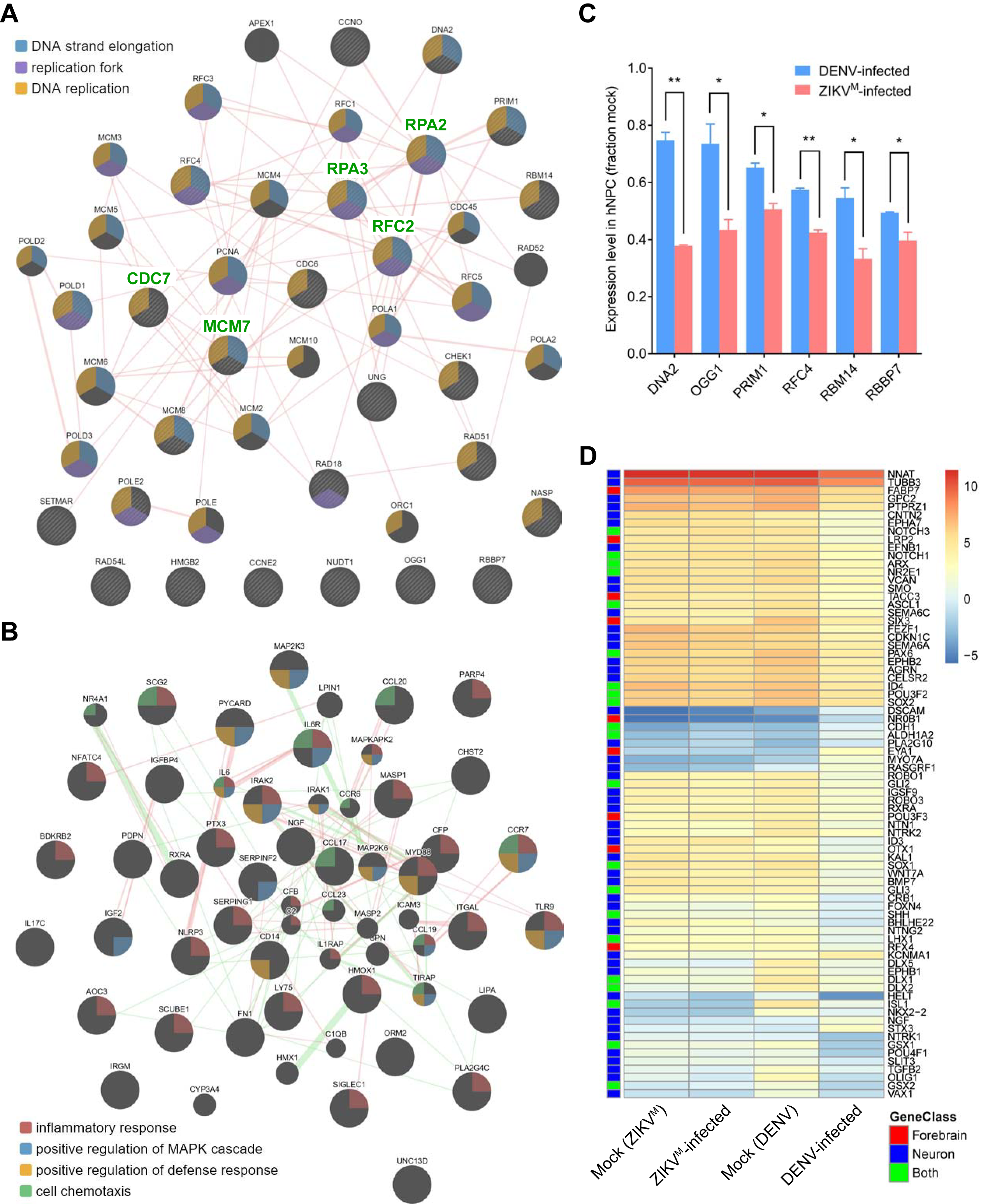
Virus-specific differentially expressed genes are enriched in DNA replication/repair, inflammatory, and neurodevelopmental pathways. (**A** and **B**) Protein-protein interaction maps highlighting (A) the protein products of 26 genes (circles with stripes) that are involved in “DNA replication” and/or “DNA repair” pathways and were only significantly downregulated in ZIKV^M^-infected hNPCs, and (B) the protein products of 37 genes (large circles) that are involved in “inflammatory response” pathway and were only significantly altered in DENV-infected hNPCs. Genes that belong to different functional groups are highlighted with colors (see legends in figures). Experimentally validated physical interactions between two proteins are indicated as pink lines. Genetic interactions between two proteins are indicated as green lines (only shown in (B)). Broader lines indicate higher confidence for each pairwise interaction. Weak interactions were removed to visually simplify the networks. (**C**) qRT-PCR results confirming the downregulated expression of DNA2, OGG1, PRIM1, RFC4, RBM4, and RBBP7 that are involved in “DNA replication” and/or “DNA repair” pathways and were significantly downregulated only in ZIKV^M^-infected hNPCs. TBP and PLA2G12A were used as internal controls. Gene expressions levels were normalized to the mock infected group. Data are represented as mean ± SD of two biological replicates (each with 3 technical replicates). *: p ≤ 0.05; **: p ≤ 0.01, *t*-tests. (**D**) Heatmap showing the expression levels (log2 FPKM) of specific genes in mock and virus-infected hNPCs. Genes that were only significantly altered in DENV-infected hNPCs, and are involved in “neuron differentiation (Neuron)” or “forebrain development (Forebrain)” are labelled by blue or red boxes, respectively. Genes belonging to both GO terms are labelled by lime green boxes.

### Both Asian and African ZIKV infect hNPCs and lead to cell death

We previously demonstrated that the African strain of MR766 (ZIKV^M^) infects hNPCs and enhances caspase activation and apoptosis (18). We tested whether an Asian ZIKV isolate, FSS13025 (ZIKV^C^), can also infect hNPCs and induce cell death. The RNA genomes of these two ZIKV strains are 88.9% identical, and the amino acid sequences of their polyproteins are 96.4% identical (99.2% similar). ZIKV^C^ efficiently infected hNPCs, resulting in a 46.7 ± 3.2% (n = 6) infection rate at 64 hours post-inoculation with an MOI of 0.04, while ZIKV^M^ had a 69.8 ± 9.5% (n = 6) infection rate after the same period of time with an MOI of 0.02 (Figure 3A). In addition, infection by ZIKV^C^ also caused increased cell death under these conditions, doubling the percentage of activated Caspase-3^+^ cells (5.2 ± 1.7% (n = 6) activated Caspase-3^+^ for ZIKV^C^-infected, 2.5 ± 0.6% (n = 5) in mock, and 9.6 ± 3.6% (n = 6) for ZIKV^M^-infected). Both strains of ZIKV reduced BrdU incorporation by infected hNPCs to a similar degree (data not shown). Thus, both Asian and African ZIKV infect hNPCs, stunt cell proliferation, and lead to cell death.

**Figure 3.**
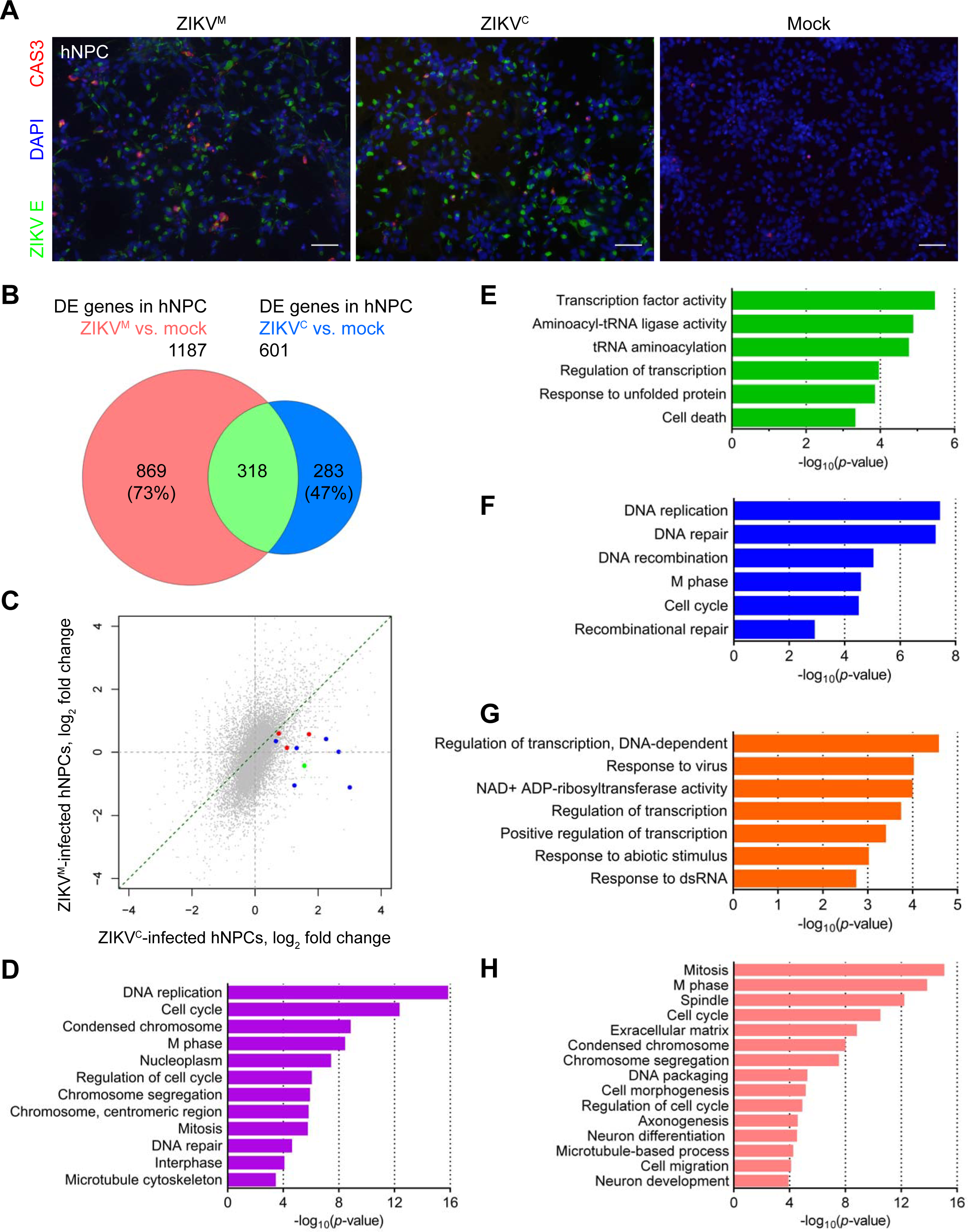
Distinct gene expression changes caused by Asian ZIKVC and African ZIKV^M^ in hNPCs. (**A**) Sample images of immunostaining of forebrain-specific hNPCs 64 hours after infection with ZIKV^M^ at a MOI of 0.02 and ZIKV^C^ at a MOI of 0.04 showing increased cell death. ZIKV envelop protein (ZIKVE; green)-positive cells were infected by ZIKV. Cleaved-caspase-3 (CAS3; red)-positive cells were undergoing cell death. DAPI (blue) was used to stain nuclei. Scale bars, 50 μm. (**B**) Weighted 2-D Venn diagram showing overlap (pale green) of differentially expressed genes identified by RNA-seq from ZIKV^M^- (salmon, MOI = 0.02) and ZIKV^C^- (royal blue, MOI = 0.04) infected hNPCs at 64 hours postinfection. (**C**) Scatter plot of global differences in transcriptome changes between ZIKV^M^- and ZIKV^C^-infected hNPCs. Log_2_ fold change of FPKM from RNA-seq data are plotted. Genes that were only significantly upregulated in ZIKV^C^-infected hNPCs, and are involved in “response to virus” and “response to dsRNA” pathways are highlighted in blue and red, respectively. The gene (STAT1) belonging to both GO terms is highlighted in green. (**D**, **E**, **F**, **G**, and **H**) GO analyses showing biological pathways, cellular compartments, and molecular functions enriched in specific sets of genes from ZIKV-infected hNPCs. Colored magenta, genes significantly downregulated in both ZIKV^C^- and ZIKV^M^-infected cells (D); lime green: genes significantly upregulated in both ZIKV^C^- and ZIKV^M^-infected cells (E); blue, genes significantly downregulated only in ZIKV^C^-infected cells (F); orange, genes significantly upregulated only in ZIKV^C^-infected cells (G); and salmon, genes significantly altered only in ZIKV^M^-infected cells (H). See also Figure S2, and Table S2.

### Asian ZIKV^C^, but not African ZIKV^M^, upregulates TP53 and viral response genes in hNPCs

To determine the molecular signatures associated with each strain of ZIKV, we compared the gene expression profiles of hNPCs infected with ZIKV^C^ and ZIKV^M^ (Figure S2 and Table S2). Overall, the gene expression changes caused by both strains of ZIKV displayed significant overlap (p < 0.0001; Chi-square test with Yates correction) (Figure 3B). In addition, transcriptome-wide analysis of fold changes in gene expression indicated that ZIKV^C^-infected hNPCs exhibit less prominent transcriptome changes than ZIKV^M^-infected hNPCs under our infection conditions (with a higher MOI but a lower infection rate for ZIKV^C^) (Figure 3C). For genes that were differentially expressed in at least one ZIKV infection group, 83.9% of them had, on average, 2.02-fold larger absolute fold changes of expression in ZIKV^M^-infected cells than in ZIKV^C^-infected cells, while only 16.1% of them had, on average, 1.47-fold larger absolute fold changes of expression in ZIKV^C^-infected cells than in ZIKV^M^-infected cells (Table S2). Genes consistently downregulated upon infection by these two strains are mainly involved in DNA replication (22 genes), cell cycle (34 genes), and DNA repair (13 genes) (Figure 3D), while the upregulated genes are associated with responses to unfolded protein and cell death (Figure 3E). Compared with ZIKV^M^, ZIKV^C^, which is more closely related to current epidemic strains, led to the dysregulation of 10 and 13 additional genes involved in DNA replication and DNA repair, respectively (Figure 3F), despite the fact that the overall number of genes with altered expression is lower in ZIKV^C^-infected cells (Figure 3B and S2). For example, TP53 (tumor suppressor protein p53) was significantly upregulated by 80% (p = 7.5E-8) in ZIKV^C^-infected hNPCs, but it was only increased by 4% (p = 0.73) in ZIKV^M^-infected hNPCs, according to RNA-seq data (Figure 4A). These differences have been further validated by qRT-PCR (Figure 4B). The differential upregulation of TP53 prompted us to test p53 inhibitors in ZIKV-induced apoptosis in hNPCs. After incubation with ZIKV (MOI = 5), caspase-3 activity in hNPCs increased dramatically and the p53 inhibitors pifithrin-α, *p*-nitro-pifithrin-α, and pifithrin-μ dose-dependently inhibited the activation of caspase-3 by both ZIKV strains (Figure 4C and 4D). However, in comparison to ZIKV^M^ infection, these p53 inhibitors clearly exhibited higher potency in rescuing ZIKV^C^-induced apoptosis in hNPCs, with 46 μM of pifithrin-α and 15 μM of *p*-nitro-pifithrin-α completely reversing the effects of ZIKV^C^-induced caspase-3 activation (Figure 4C and 4D). Neither nutlin-3B, an activator of p53 (39), nor *p*-nitro-pifithrin-α exhibited direct effects on ZIKV production (Figure S3). In addition, GO terms “response to virus” and “response to dsRNA” were significantly enriched in genes upregulated only in ZIKV^C^-infected cells (Figure 3G and 4E), but not in upregulated genes shared between ZIKV^C^-infected and ZIKV^M^-infected cells (Figure 3E). Examples of these ZIKV^C^-specific upregulated genes are interferon-stimulated genes ISG15 and MX1; STAT1, IRF9, CXCL10, OAS1, IFIT2, G1P2 and G1P3 in Type II interferon signaling (adjusted p = 4.3E-6); and STAT1, CXCL10, NFKB2 and NFKBIA in Toll-like receptor signaling pathway (adjusted p = 0.0081). Upregulated expression of ISG15 and STAT1 in ZIKV^C^-infected hNPCs was validated by qRT-PCR (Figure 4B). ZIKV^C^-specific altered genes were also significantly enriched in TNFα signaling pathway (NEKBIA, NFKBIE, REL, NFKB2 and PTPRCAP; adjusted p = 0.0067). Consistent with the Asian strain-specific antiviral response in the infected hNPCs, quantitation of ZIKV gRNA in hNPCs at 64 hours post-infection revealed a much lower viral load in ZIKV^C^-infected hNPCs than in ZIKV^M^-infected hNPCs, despite similar initial MOIs (Figure 4F). Furthermore, ZIKV^M^-specific DEGs were highly enriched in mitosis and cell cycle-related pathways, and moderately enriched for the GO term “neuron differentiation”. Among the 35 DEGs belonging to the GO term “neuron differentiation”, 21 and 28 also belonged to “axonogenesis” and “neuron development”, respectively (Figure 3H). These observations suggest that different ZIKV strains can impact common pathways involved in the regulation of hNPC proliferation, but that there is strain-specific gene regulation as well.

**Figure 4.**
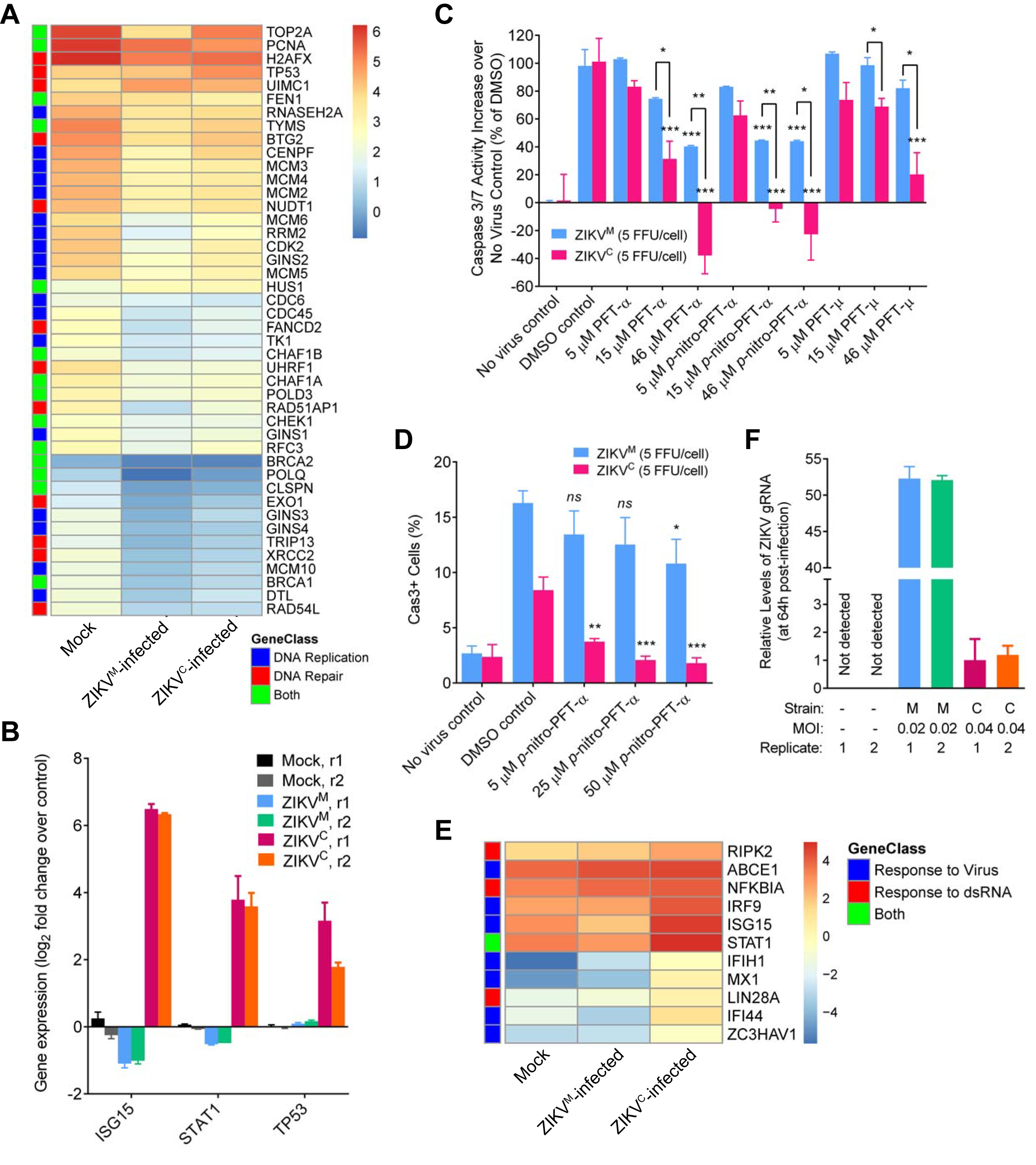
Asian ZIKV^C^, but not African ZIKV^M^, induces the upregulation of TP53 and viral response genes in infected hNPCs. (**A**) Heatmap showing the expression levels (log2 FPKM) of specific genes in mock and ZIKV-infected hNPCs. Genes that were significantly altered in ZIKV^C^-infected hNPCs, and are involved in “DNA replication” and “DNA repair” pathways are labeled by blue and red boxes, respectively. Genes belonging to both GO terms are labeled by green boxes. (**B**) qRT-PCR results confirming the upregulated expression of ISG15 and STAT1 that are involved in “response to virus” pathway and were significantly upregulated only in ZIKV^C^-infected hNPCs, and TP53 that was specifically altered in ZIKV^C^-infected hNPCs. TBP was used as an internal control. Gene expressions levels were normalized to the mock infected group. Data are represented as mean ± SD of two technique replicates. r1 and r2: biological replicates 1 and 2. (**C**) Inhibition of ZIKV-induced caspase-3 activity in hNPCs by p53 inhibitors. Caspase-3 activity in hNPCs increased after 6-hour incubation with ZIKV^M^ or ZIKV^C^. All three p53 inhibitors (pifithrin (PFT)-α, PFT-μ, and *p*-nitro-PFT-α) dose-dependently reversed the ZIKV-induced caspase-3 response, with higher potency in ZIKV^C^-infected cells than in ZIKV^M^-infected cells. Data are represented as mean ± SD (n = 2). Inhibitor-treated groups were compared with the DMSO-treated group, and ZIKV^M^-infected groups were compared with ZIKV^C^-infected groups. *: p ≤ 0.05; **: p ≤ 0.01; ***: p ≤ 0.001; p-values larger than 0.05 are not shown, *t*-tests. (**D**) Quantitation of Cas3+ cells in ZIKV-infected and p53 inhibitor-treated hNPCs by immunocytochemistry. Cas3+ cells increased in ZIKV-infected (MOI = 5) hNPCs at 72-hour post-infection. A p53 inhibitor, *p*-nitro-PFT-α, dose-dependently decreased the percentage of Cas3+ cells in ZIKV^C^-infected hNPCs, and completely reversed the effect of ZIKV^C^ infection at concentrations of 25 μM and above. However, the potency of *p*-nitro-PFT-α was found to be much lower in ZIKV^M^-infected hNPCs. Data are represented as mean ± SD (n = 3). Inhibitor-treated groups were compared with the DMSO-treated group. *ns*: not significant, p > 0.05; *: p ≤ 0.05; **: p ≤ 0.01; ***: p ≤ 0.001; t-tests. (**E**) Heatmap showing the expression levels (log2 FPKM) of specific genes in mock and ZIKV-infected hNPCs. Genes that were only significantly upregulated in ZIKV^C^-infected hNPCs, and are involved in “response to virus” and “response to dsRNA” pathways are labeled by blue and red boxes, respectively. The gene (STAT1) belonging to both GO terms is labeled by a green box. (**F**) qRT-PCR results showing the relative levels of ZIKV gRNA in hNPCs at 64 hours post infection. TBP was used as an internal control. The levels of ZIKV gRNA were normalized to the ZIKV^C^-infected hNPCs. Data are represented as mean ± SD of two technique replicates.

## DISCUSSION

A recent increase in the rate of microcephaly has been associated with an outbreak of ZIKV, a Flavivirus mainly transmitted by mosquitoes and through sexual transmission (3–5). Recent evidence has shown that ZIKV is able to infect human neural progenitor cells, neurospheres and brain organoids (18–20). But how the different target cell types in the nervous system respond to ZIKV infection and contribute to brain developmental defects is unknown. Here we profiled gene expression in hNPCs exposed to Asian ZIKV^C^, African ZIKV^M^, and dengue virus (DENV), and identified virus- and strain-specific alterations of gene expression in hNPCs. The gene expression profiling datasets we present here are the first systematic analyses of transcriptomic changes induced by different strains of ZIKV in its target cell types. Historically, gene expression profiling has been used to elucidate biological pathways and underlying mechanisms to reveal previously unknown subtypes of a disease, and to predict disease prognosis (40). Utilization of our datasets may lead to a better understanding of the molecular pathogenesis of ZIKV infection-induced brain abnormalities, and potentially facilitate identification of biomarker(s) for disease diagnosis and treatment. Moreover, analysis of these transcriptomic profiles in the context of the established Connectivity Map, which links gene expression changes to specific small molecules, could reveal candidate small molecules to reverse or prevent the biological responses induced by ZIKV infection, which could have therapeutic benefits for ZIKV-infected individuals (40).

Many flaviviruses have broad cellular and species tropism and multiple factors besides viral susceptibility may contribute to pathogenic outcomes, including specific cellular responses and tissue access. DENV, a flavivirus with genetic similarities to ZIKV, has been shown to infect cells of several lineages but hematopoietic cells play an essential role in its pathogenesis (41). Recently published work, and our results presented here, suggest that dengue virus 2 (DENV2) could infect hNPCs efficiently; however, it does not appear to significantly impact the proliferation of hNPCs (unpublished data), the morphology of neurospheres (19), or the growth of brain organoids (19). Our transcriptome profiling analyses suggest that, although the acute exposure to DENV2 could influence the expression of the genes involved in DNA replication and DNA repair, ZIKV infection results in a much more targeted impact as reflected by a higher percentage of genes with altered expression in these pathways. Interestingly, DENV infection-induced changes in gene expression were involved in inflammatory response and Wnt signaling. These findings suggest that DENV2 and ZIKV target distinct biological pathways.

Genetic and phylogenetic analyses of ZIKV have identified two main ZIKV lineages, African and Asian (42). Strains implicated in the recent epidemic have been traced to the Asian lineage (13). Although ZIKV has been circulating throughout Africa and Asia since 1947, ZIKV infections were not found to be associated with significant human pathology until now. As a first step to gain a better understanding of ZIKV pathogenesis, we compared the impact of infection by ZIKV of both African and Asian lineages on hNPCs. Both strains could infect hNPCs efficiently and led to increased cell death. Gene expression analyses suggest significant overlap of genes with altered expression in infected hNPCs between African and Asian strains, indicating that the mode of action of these two viral strains is similar once exposed to the fetal brain. Interestingly, the Asian strain that we examined, which is more closely related to the strain implicated in the current epidemic, induced specific dysregulation of DNA replication and repair genes (e.g., TP53), and the upregulation of viral response genes involved in interferon response, Type II interferon signaling, Toll-like receptor signaling and TNFα signaling pathways in hNPCs.

We have shown that three p53 inhibitors dose-dependently inhibited the activation of caspase-3 in hNPCs induced by the two ZIKV strains, with significantly higher potency in ZIKV^C^-infected cells than in in ZIKV^M^-infected cells. This finding is consistent with the strain-specific upregulation of TP53 upon infection of hNPCs by the Asian strain, as revealed by our RNA-seq and qRT-PCR data. Pifithrin-α reversibly inhibits p53-dependent transactivation, and also protects cells from p53-mediated apoptosis induced by various stimuli (43). *p*-Nitro-pifithrin-α, a cell-permeable analog of pifithrin-α, is 10 times more potent than pifithrin-α in inhibiting p53 activity (44). Indeed, *p*-nitro-pifithrin-α displayed higher potency in reversing caspase-3 activation for both ZIKV^M^- and ZIKV^C^-infected hNPCs (Figure 4C). Pifithrin-μ inhibits p53-mediated apoptosis by disrupting its binding to Bcl-xL and Bcl-2 in mitochondria but does not affect p53-dependent transactivation (45). Interestingly, at concentrations up to 46 μM, pifithrin-μ significantly inhibited ZIKV^C^-induced caspase-3 activation but not ZIKV^M^-induced activation (Figure 4C). Together, these findings suggest that p53 plays pivotal roles in the Asian strain-induced apoptosis in hNPCs, and p53 inhibitors could provide protection during ZIKV infection in the central nervous system (CNS).

Accumulation of viral structural proteins has been shown to induce apoptosis in infected cells via upregulation of p53. Two examples are the *N*-terminus of the capsid protein (Cp) of Rubella virus (RV) in the *Togaviridae* family (46,47), and the *C*-terminus of West Nile virus (WNV) Cp (48–50). It is tempting to speculate that the ability of elevating p53 expression resides in the Cp of ZIKV^C^, given the 12-amino acid (a.a.) difference between the full-length ZIKV^M^ Cp and ZIKV^C^ Cp, and especially the 4-a.a. difference within the last 22 residues in the *C*-terminus (R101K, A106T, I110V and 1113V). However, caution should be exercised, for two reasons. First, the mRNA level of p53 was upregulated by ZIKV^C^ infection in hNPCs (Figure 4A and 4B), but the mRNA levels of p53 were unchanged during both WNV and RV infection (46,50). Secondly, phosphorylation of WNV Cp at residues near Ser83 or within Ser99 and Thr100 by protein kinase C is essential for WNV Cp to sequester HDM2 and stabilize p53 (51). But neither of the phosphorylation sites can be found in ZIKV Cp. Specifically, in both ZIKV^M^ Cp and ZIKV^C^ Cp, Ser83 in WNV Cp is substituted by Lys83, and equivalent residues of Ser99 and Thr100 in WNV Cp are absent. Therefore, the upregulation of TP53 in hNPCs by ZIKV^C^ and the elevation of p53 protein by WNV are most likely through different mechanisms.

In summary, our genome-wide gene expression analyses reveal virus- and strain-specific molecular signatures associated with ZIKV infection of human cortical neural progenitors. These data will help guide future investigations of ZIKV-host interactions and help illuminate neurovirulence determinants of ZIKV in patients.

## SUPPLEMENTARY DATA

Supplementary Data are available at NAR Online.

## ACKNOWLEDGEMENT

We thank Dr. Robert Tesh at UTMB and the World Reference Center for Emerging Viruses and Arboviruses (WRCEVA) for FSS 13025 isolate of ZIKV, Lihong Liu and Yuan Cai of the Ming and Song labs for technical assistance. In addition, we would like to thanks C. Strauss for critical reading of the manuscript, Weining Tang at Omega Bioservices (Omega Bio-tek, Inc) and Mike Zwick and Ben Isett at the Emory Integrated Genomics Core (EIGC) for assistance with high-throughput sequencing.

